# Markerless tracking suggests a tactile sensing role for forelegs of *Dolomedes* spiders during locomotion

**DOI:** 10.1101/398479

**Authors:** Kiri F. Pullar, Michael G. Paulin

## Abstract

**Summary statement:** We developed a machine vision technique for markerless tracking of locomotion in the spider *Dolomedes aquaticus.* Gait analysis suggests that each pair of legs plays a specific role in locomotion.

**Abstract:** Because of their rigid exoskeleton with relatively simple joint mechanics, arthropods can provide useful models for studying the sensory-neural and mechanical design principles of agile animal locomotion. Gait analysis usually requires attaching markers or manually identifying reference points in video frames, which can be time consuming and inaccurate, especially with small animals. Here we describe a markerless motion capture technique and its application to gait analysis in the New Zealand semi-aquatic hunting spider, *Dolomedes aquaticus*. Our machine vision approach uses a model of the spider’s skeleton to infer the location of the centre of mass and the configuration of the skeleton in successive video frames. We found that stride length and frequency are correlated with running speed. Inter-limb coordination during the gait cycle suggests that different legs have specialized roles in locomotion. Phase relationships among the six hindmost legs exhibit an alternating tripod gait, as in hexapod insects. The middle two leg pairs appear to be primarily responsible for generating thrust, while the hind legs contribute more to stability. The front legs are not phase-coupled to the other legs and appear to be used as tactile probes during locomotion. Our machine vision approach has the potential to automate arthropod gait analysis, making it faster and easier. Our results indicate how specialization of limb function may contribute to locomotor efficiency and agility of a specialized hunting spider, and how arthropod design principles may contribute to developing efficient, agile legged robots.

## Introduction

Arthropods have proven favourable subjects for studies of locomotion for over a century (Bowerman, 1977). Apart from being of interest in their own right, there are several reasons why investigations into the neural, morphological and mechanical mechanisms underlying arthropod locomotion have broader appeal. Firstly, locomotion is essential for life in the majority of organisms, playing an important role in interactions between mates, predators and prey. Secondly, repeated stereotyped behaviour can be produced with relative ease which allows for descriptive, quantitative and neurophysiological analyses (Bowerman, 1977). Finally, the characteristic rigid exoskeleton and jointed limbs of arthropods can be represented as a rigid body linkage, in which most joints only have a single degree of freedom. Despite major morphological and physiological differences fundamental patterns appear to exist in locomotion, thus generalizations in arthropods can be extrapolated to make predictions in systems of greater complexity e.g. vertebrates and robots (Dickinson et al., 2000).

Although arachnids are one of the largest arthropod groups, study of their locomotory behaviour has been somewhat neglected compared to insects and crustaceans. While the collection of studies is smaller in magnitude they still amount to a reasonably comprehensive analysis of various aspects of locomotion: behavioural descriptions of normal locomotion (Biancardi et al., 2011; Bowerman, 1975a; Shamble et al., 2017; Spagna et al., 2011; Ward and Humphreys, 1981; Weihmann, 2013; Wilson, 1967; turning - Land, 1972; water surface - Shultz, 1987; temperature dependence - Booster et al., 2015; prey strikes - Zeng and Crews, 2018), behavioural descriptions of experimentally modified animals (load carrying - Moffett and Doell, 1980; legs removed or immobilized - Amaya et al., 2001; Apontes and Brown, 2005; Bowerman, 1975b; Gerald et al., 2017; Wilson, 1967), joint kinematics (Blickhan and Barth, 1985; Booster et al., 2015), neurophysiological examinations (Bowerman, 1976; Bowerman and Burrows, 1980; Fourtner and Sherman, 1973; Maier et al., 1987), anatomy of locomotor appendages (Blickhan and Barth, 1985; Bowerman and Burrows, 1980; Dillon, 1952; Fourtner and Sherman, 1973; Parry, 1957; Reußenzehn, 2008; Reußenzehn, 2010), biomechanics (Anderson and Prestwich, 1975; Blickhan, 1986; Blickhan and Barth, 1985; Ellis, 1944; Parry and Brown, 1959a; Reußenzehn, 2010; Sensenig and Shultz, 2006; Shultz, 1991; Siebert et al., 2010; Wilson, 1970; Wilson and Bullock, 1973) and dynamics of locomotion (Biancardi et al., 2011; Blickhan and Barth, 1985; Sensenig and Shultz, 2006; Spagna et al., 2007; Spagna et al., 2011; Weihmann, 2013; jumping - Parry and Brown, 1959b; prey strikes - Zeng and Crews, 2018).

The present study focuses on the stepping patterns and kinematics involved with normal spider locomotion. Similar analyses mentioned above generally used one of two approaches. Marker-based techniques, which involve the placement of markers (e.g. paint, nail polish, paper or tape) onto the centre of a joint or tip of a limb. The marker is then tracked in video frames, usually through the aid of commercial software packages. Alternatively, the researcher manually digitized points of interest in each frame through the use of a stop-motion projector. Both approaches are laborious in nature and require careful attention, the first at the beginning of the experiment to ensure the correct placement of markers and the second during processing to accurately locate points of interest. These problems are compounded when working with smaller animals, where positioning of markers, sensors or points of interest is more critical in an accurate description of locomotor kinematics (Gambaretto and Corazza, 2009). This limits the researcher’s ability to examine locomotion, as markers can interfere with the animals normal movement and behaviour (Bender et al., 2010). Additionally, although maker-based approaches are widely accepted, they come with intrinsic problems e.g. ghost markers, occlusions of markers, markers can move or become completely removed during locomotion and often require specialized lab environments and lighting conditions. To address the concerns above, we developed semi-automated model-based tracking technique that can capture the 2D motion of the body and leg segments from a high-speed video sequence.

Advances in imaging and computer vision are enabling increasingly complete records of animal motion (Brown and Bivort, 2018). Robust approaches for tracking and analysis of body movements have been used to examine locomotion in a variety of animals. Dimensionality reduction methods can be used to identify pixels of video frames that change during animal behaviour and match spatio-temporal image features to exemplar postures, behaviours, or gaits (Berman et al., 2014; Favreau et al., 2006; Gibson et al., 2003; Klibaite et al., 2017). Pixel-based representations can be applied to animals regardless of their shape or limb configuration (Brown and Bivort, 2018), however accuracy of tracking is often improved by utilizing information about the shape, appearance, and/or kinematic structure of subject being tracked (Moeslund et al., 2006).

Model-based approaches incorporate information about the subject to estimate its most likely pose given the image sequences (Brown and Bivort, 2018). A motion model was developed to track wallaby locomotion, estimating the pose of a kinematic chain from frame to frame (Bregler et al., 2004). The study originally intended to examine locomotion via a marker-based approach, however it became clear that the locations of makers could not be accurately tracked due to soft tissue deformations (Bregler et al., 2004). Models can be extended to deform not only a skeleton (i.e. the kinematic chain), but also the non-rigid deformation of the 3D surface (i.e. clothing, skin or fur). Several studies of dog locomotion have used probabilistic approaches to match a body shape model to the corresponding silhouette in image frames (Gall et al., 2009; Gambaretto and Corazza, 2009). Few studies have implemented similarly robust methods with the precision required for automated, markerless, tracking of arthropod bodies and appendages (Uhlmann et al., 2017).

An investigation of flight kinematics in *Drosophila*, used an accurate geometric fly model, scaled motion dynamics and image registration to capture body and wing rotations (Fontaine et al., 2009). While the kinematic chain in this case required relatively few degrees of freedom, its ability to track a very small animal during complex behaviours is impressive. Recently, a markerless approach has also been applied to leg segment tracking in freely walking *Drosophila* (Uhlmann et al., 2017). FlyLimbTracker used a body and leg model defined by active contours, for a semi-automatic approach which drastically reduced the user intervention required in tracking compared to manual digitization (Uhlmann et al., 2017). In addition to providing important information regarding locomotor parameters, high resolution recording and analyses also can assess the role of proprioceptive sensory inputs during coordinated leg movements (Gowda et al., 2018).

In the present study we develop a semi-automated, model-based, markerless approach for tracking freely moving spiders. We focused on providing a detailed analysis of locomotory kinematics without the need for a sophisticated lab setup, expensive commercial software or manual digitization of joint location in each frame. Based on the 2D position of joints in each frame we extract data on the trajectory and coordination of each of the four pairs of legs and compare the results to existing literature.

## Materials and methods

### Animals

The experimental procedure consisted of filming the locomotor patterns of unrestrained *Dolomedes aquaticus* Goyen. This species is commonly found near water on or under surrounding objects such as plants, stones and pieces of wood. *D. aquaticus* sense insects and small vertebrates both on water and land by detecting vibrations using the two front pairs of legs (Forster and Forster, 1999). They then chase down their prey with rapid bursts of speed. Only relatively large, adult, female spiders possessing all eight legs were used during filming (mass = 1.194 g ± 0.376; mean ± S.D.).

### Experimental approach

Behavioural arenas consisted of a custom made 40 cm × 10 cm × 2 cm glass arenas; this size was selected as large enough for free locomotion but small enough to encourage spiders to run in a relatively straight line. The arena was lit from below using three 10 × 10 arrays of ultra-bright LEDs. In order to achieve a more uniform light intensity, layers of frosted glass and frosted paper were used to scatter the light from the LEDs. A piece of cardboard covered the last 5 cm at one end to the spider to act as a refuge from the bright light. A 1 cm grid was printed onto an overhead projector transparency overlaid on the arena. The video camera was positioned in the middle, above the arena, to allow filming of the spider from a dorsal viewpoint (Fig. 1A). Trials were recorded using a high-speed video camera (125 fps, 1/625 s shutter speed, 1280 × 1024 pixels with a Troubleshooter 1000 HR, Fastec Imaging, San Diego, USA).

**Fig. 1.**
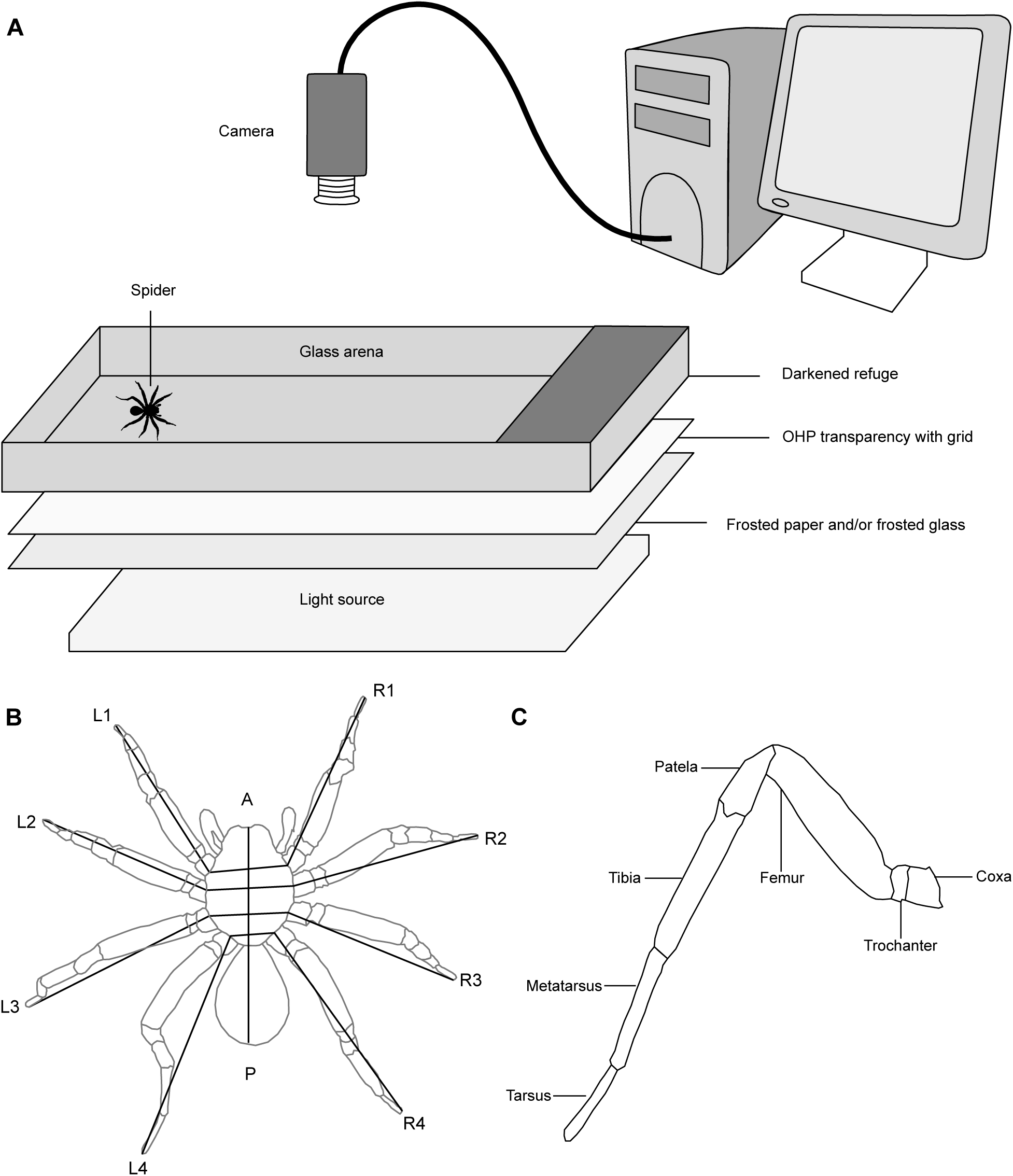
Experimental setup and study animal. (A) Experimental setup consists of a glass arena with a darkened refuge at one end. The arena is lit by LED arrays scattered using frosted paper and/or frosted glass. A video camera films the spider from above. The camera was attached to a computer, which was used for image capture, storage and analysis. For further details see text. (B) Locomotor appendages of the spider. Leg movements were based on the angle delimited by the tarsus, the body-Coxa joint of a given leg and the body-Coxa joint of the contralateral leg. This angle was equal to 0 when the leg was perpendicular to the cephalo-caudal axis of the body. Its value increased when the leg moved forwards towards the AEP and decreased when the leg moved backwards towards the PEP. (C) Morphology of the spider leg showing the segments involved in locomotion.

Spiders were occasionally stimulated to move from one end of the arena to the other by either prodding them with tweezers or gently blowing on them. The resulting locomotion did not appear to differ from normal running in unstimulated animals. Each animal was run no more than five consecutive times. Short rest periods were given between each trial while data was transferred from the camera to the computer and converted into HD resolution avi files using the camera software (CamLink, Fastec Imaging, San Diego, USA). Trials where the animal did not run in a straight line or at a near constant speed were rejected. The animals were not observed slipping on the glass surface and did not appear to have any difficulty running. Starting and stopping sequences were not used in analysis.

The locomotor patters of 15 spiders were filmed and their kinematic parameters analysed. Of the many sequences filmed, only a small portion were selected for analysis on the basis of the requirements of constant velocity and straight-line locomotion (N=29). The number of sequences is small due to the small portion of trails that were suitable for analysis as well as the lengthy process involved in pose estimation.

### Pose estimation algorithm

This section will briefly describe the algorithm used to estimate the pose of the subject in each frame. This is done using the PoseCut algorithm, which involves energy minimization based on a Conditional Random Field (CRF) in order to estimate the pose of the subject in each frame of a video sequence, as well as segment each frame into subject and background (Bray et al., 2006). The pose estimation code was based around adapting the techniques used by Rasmus Jensen for analysis of human gait using a time of flight camera. The pose estimation algorithm is briefly described below, for further detail see Jensen et al. (2009).

A frame of the sequence (Fig. 2A) can be treated as set of discrete random variables ***y*** *= y*_*1*_*, y*_*2*_*,…, y*_*n*,_ where each *y*_*i*_ represents the intensity of the pixel *i* of an image containing *n* pixels. The solution to our image segmentation problem becomes finding the value of the vector ***x*** *= x*_*1*_*, x*_*2*_*,…, x*_*n*_ where each *x*_*i*_ represents the label assigning pixel *i* to an object. Each *x*_*i*_ takes its value from the label set *L = l*_*1*_*, l*_*2*_*,…, l*_*m*_, which in our case is a binary problem with labels consisting only of ‘background’ and ‘subject’.

**Fig. 2.**
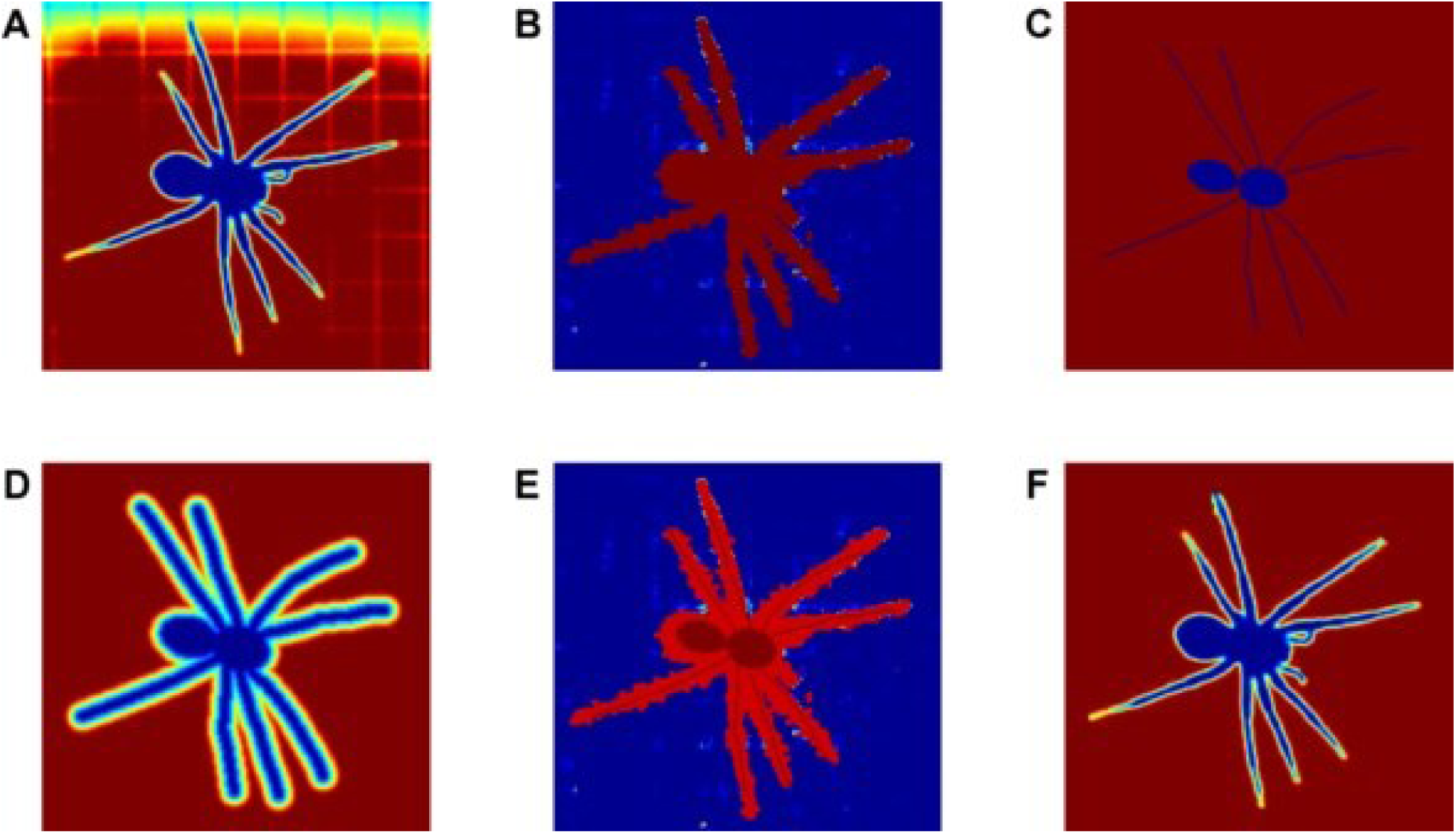
Different terms in our image segmentation and pose estimation algorithm. (A) Original image. (B) Likelihood of pixels being labelled background. (C) The stick spider model after optimization of its 2D pose. (D) The shape prior (distance transform) corresponding to the optimal pose of the stick spider. (E) Likelihood of pixels being labelled background after incorporation of the shape prior. (F) The segmentation result obtained by the algorithm which is the maximum a posteriori solution of the energy of the pose specific CRF.

The first term of the energy function is a likelihood term, which is based on the intensity distribution of background pixels (Fig. 2B). This term is calculated as the negative log likelihood:

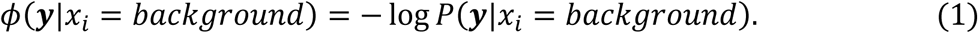

Assuming the background pixels have distribution with mean, *µ*_*B*_, and standard deviation, *σ*_*B*_, the probability that observed data, ***y***, belongs to the background is given by:

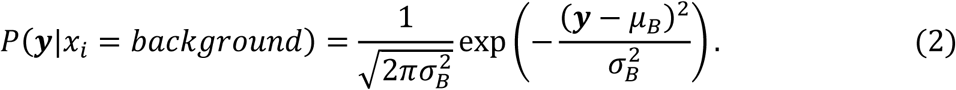

By taking the negative log of the probability function and discarding the normalization constant, this becomes:

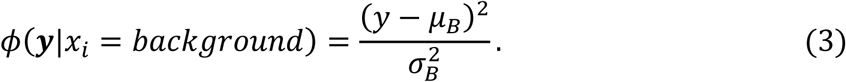

The beginning of each sequence contained at least 5 frames of background before the subject entered the arena. Thus, the intensity distribution is estimated by simply calculating the mean and standard deviation of each pixel in known background frames.

The second term of the energy function is the smoothness prior, which specifies neighbouring pixels will have a higher probability of having the same label. The prior takes the form of a generalized Potts model (Bray et al., 2006):

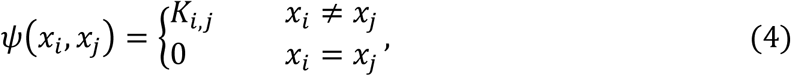

where *K*_*i,j*_ is a term that penalizes adjacent neighbours which have different labels. Because we are dealing with the binary situation of segmenting between background and subject the problem is referred to as an Ising model (Greig et al., 1989).

Additionally, it is assumed that neighbouring pixels with the same label will have similar intensity value. A contrast term is incorporated by increasing the cost within the Ising model proportionally to the similarity in intensity of the corresponding pixels. Thus, penalizing neighbours with similar intensity values which have differing labels:

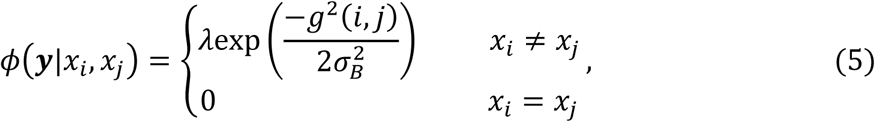

where the parameter *λ* controls the magnitude of the penalty and *g*^*2*^(*i, j*) is the gradient, which is approximated via convolution of the image with gradient filters in both in horizontal and vertical directions.

The final term in the energy formulation is a shape prior, this looks for an object with known shape and plays an important role in pose estimation. An articulated stick spider model is used to generate a pose-specific shape prior on the segmentation. The stick spider is based on measurements of the length and maximum joint angles of each leg segment for *D. aquaticus* (Reußenzehn, 2008). Given an estimate of the location and shape of the subject, the pose-specific prior is constructed so that pixels falling close to the shape have a higher probability of being labelled ‘subject’ while those falling further away have a higher probability of being labelled ‘background’.

The cost of the shape prior is defined as:

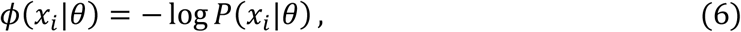

where *θ* contains the configuration of the stick spider (position, orientation and leg span). The probability *P*(*x*_*i*_|*θ*) of *x* being labelled as subject or background is defined as:

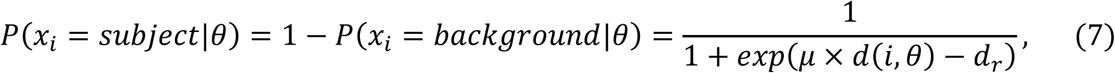

where *d*(*i,θ*) the distance of pixel *i* from the shape defined by *θ*, *d*_*r*_ is the thickness of the shape and μ contains the magnitude of the cost for points outside of the shape. To calculate the distance of each pixel from the shape, the rasterized model (Fig. 2C) undergoes the Euclidean distance transform, resulting in a distance map where pixels that are part of the shape get value zero, those adjacent to the shape get value 1 and so on increasing in magnitude (Fig. 2D).

The combination of the terms described above is the energy formulation (Fig. 2E) that is used both to segment the frames and estimate the pose of the subject (Fig. 2F):

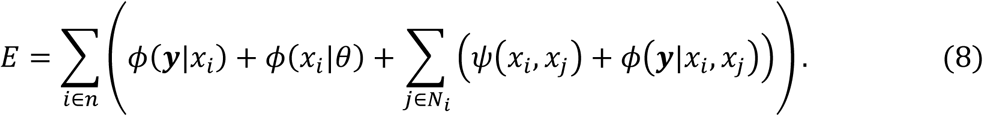

This energy corresponding to the configuration of the CRF is minimized using the graph cuts approach (Boykov and Kolmogorov, 2004). Each frame requires an initial guess which is optimized to find the estimated pose. In this case the algorithm begins with the pose of the subject in the previous frame updated under the assumption of smooth motion. The assumption of constant velocity provides an initial guess.

The only frame requiring manual initialization is the first frame used for tracking. A Graphical User Interface (GUI) was created to allow the user to set the starting pose of the spider in a frame of each video clip and save the coordinates for batch processing later. After the video was loaded and cropped to contain the tracking region of interest, the GUI allowed the user to adjust the global position, angles and leg span of the spider. For estimating the angles, it generally suffices to only set the body-coxa joint for each leg and global rotation. The body and segment lengths defined in the shape model are used to create a model of the spider that has been adjusted for leg span, but retains the relative proportions of each segment. The shape model is used to extract configuration values for body position and orientation and the joint angles in each frame.

### Gait analysis

Prior to analysis, all raw configuration data was smoothed using a fourth-order, zero-lag Butterworth filter with cut-off frequency uniquely determined for each of the x-y and angular positions. Residual analysis was used in each case to determine an appropriate cut-off frequency (Winter, 1990). Velocity and acceleration of the smoothed linear and angular data were approximated using the numerical method of finite differences.

The overall angular movement of the leg was approximated by the angle delimited by the tarsus, the body-coxa joint and the body-coxa joint of the contralateral leg (Fig. 1B). Angles are reported with zero being perpendicular to the mid line of the cephalothorax. These angles increased when the leg moved forwards, reaching a maximum corresponding to the anterior extreme position (AEP) and decreased when the leg moved backwards, reaching a minimum corresponding to the posterior extreme position (PEP). The above system is maintained for individual joints with zero being perpendicular to the mid line of the cephalothorax for the body-coxa joint and the remaining angles are reported as rotation about the line from joint to parent (Fig. 1C).

The period of the stride was defined as the time elapsing between two successive AEPs of the same leg. It could only be measured to within 8 ms because of the film speed. The period consisted of one stance phase and one swing phase. The stride frequencies were calculated by 1/stride duration and presented as the number of strides per second. Stride length was calculated by dividing the average speed by the stride frequency. The kinematic duty factor (stance duration/stride duration) was expressed as percentage of stride duration. Within a given sequence, stride duration measurements did not differ significantly between legs (p>0.05 ANOVA); however, the movements of legs 1 and 4 are more ambiguous for functional reasons i.e. dragging being included as part of protraction. Thus, unless stated otherwise, stride data is presented using leg L2 for comparisons.

Phase relationships between legs during locomotion were calculated by the time at which an AEP occurred for a given leg relative within the period of the reference leg (Jamon and Clarac, 1995):

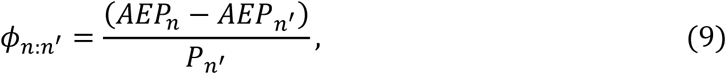

where *P*_*n’*_ is the stride period of the reference leg. Values of *Φ* less than 0.25 or greater than 0.75 represent simultaneous in-phase stepping and values of *Φ* between 0.25 and 0.75 represent alternate anti-phase stepping.

### Statistical analysis

All graphing and statistical analyses were performed in MATLAB (The MathWorks, USA). We used linear regression, one-way ANOVA and Tukey’s post hoc tests to analyse corresponding data. Descriptive statistics are presented as mean ± S.D., and the significance level is set at α=0.05. Comparisons relating kinematic parameters to speed were presented for each trial (N=29). Analysis of phase lags required that at least two full strides were completed by each leg, reducing the number of trials (N=27).

For phase lag data, each phase value was multiplied by 360 to convert from phase range (0-1) to degrees (0-360). The mean angle (phase) of leg lag was calculated in MATLAB as:

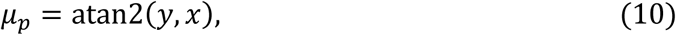

where *atan2* is a variant of the arctangent function, 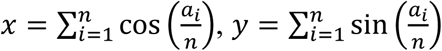, and *n* is the sample size (Fisher, 1995). This value is then divided by 360 to convert the result back to the original range. The circular standard deviation was calculated as:

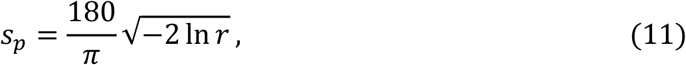

where *r* is the length of the mean vector, 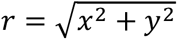 (Zar, 1996). Again, this value is then divided by 360 to convert the result back to the original range. Phase distributions were tested for uniformity of distribution using the Rayleigh test. Rayleigh’s *z*:

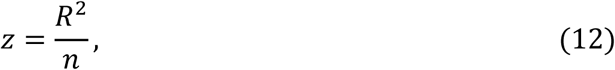

where *R* = *nr* (Zar, 1996). The p value associated with Rayleigh’s *R* is approximated by:

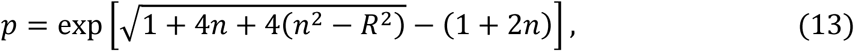

which is accurate to three decimal places for samples sizes as small as ten (Zar, 1996). To determine if any relationship existed between phase lags and speed, data was examined for angular-linear correlation. The correlation coefficient was calculated as:

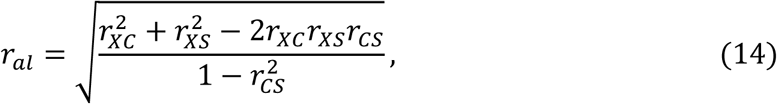

where *r*_*XC*_ is the correlation between *X* (speed) and the cosine of *a* (phase), *r*_*XS*_ is the correlation between *X* and the sine of *a* and *r*_*CS*_ is the correlation between the sine and cosine of *a* (Zar, 1996). The significance of the correlation is assessed by comparing the test statistic 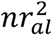 to *X*^2^distribution with 2 degrees of freedom (Zar, 1996).

## Results

The locomotion speed measured in 15 individuals during a total of 29 sequences averaged 24.807 ± 6.698 cm s^-1^ and ranged between 5.000 and 49.548 cm s^-1^. Stepping pattern diagrams (Fig. 3) for a range of speeds revealed several trends in locomotor parameters which were investigated in more detail. Stride frequency increased linearly with both absolute (y = 3.793 + 0.134x, R^2^=0.424, F_1,27_=19.848, p<0.001) and normalized speed (y = 3.174 + 0.309x, R^2^=0.580, F_1,27_=37.360, p<0.001). Stride length also increased linearly with both absolute (y = 1.694 + 0.075x, R^2^=0.464, F_1,27_=23.331, p<0.001) and normalized speed (y = 2.171 + 0.108x, R^2^=0.249, F_1,27_=8.949, p=0.006). Stride frequency was more highly correlated with normalized speed and stride length was more highly correlated with absolute speed.

**Fig. 3.**
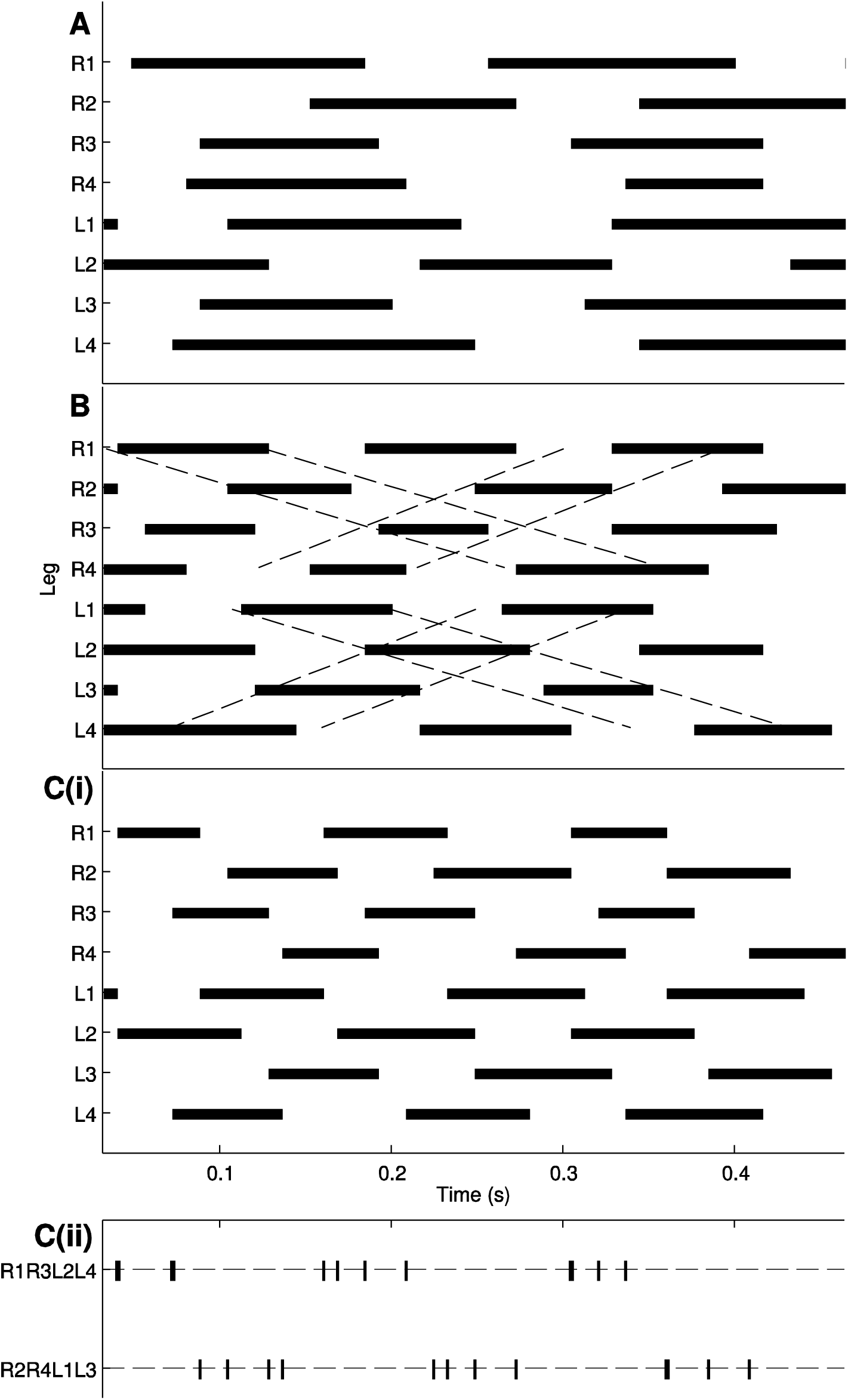
Stepping pattern diagrams from a single spider at a range of speeds. The heavy black lines represent stance phase (retraction) and the spaces swing phase (protraction). (A) Average speed 15.963 cm s^-1^, stride frequency 5.142 Hz and stride length 3.116 cm. (B) Average speed 24.286 cm s^-1^, stride frequency 6.632 Hz and stride length 3.691 cm. Dashed lines illustrate apparent forward or backward metachronal waves. (C) Average speed 33.306 cm s^-1^, stride frequency 8.403 Hz and stride length 3.997 cm. (C (ii)) The data is rearranged to emphasise functional leg groups. Onset of retraction is represented by a vertical line; the alternating tetrapod is more evident. The scale is located at the bottom of C (i).

The positive trend between stride frequency and speed suggests the lengths of the protraction and/or retraction phases may also have a relationship with speed and stride frequency. Both protraction and retraction periods decreased with absolute (protraction: y = 0.105 – 0.001x, R^2^=0.284, F_1,27_=10.703, p=0.003; retraction: y = 0.121 - 0.002x, R^2^=0.323, F_1,27_=13.034, p=0.001) and normalized speed (protraction: y = 0.111 - 0.003x, R^2^=0.384, F_1,27_=16.808, p<0.001; retraction: y = 0.128 – 0.004x, R^2^=0.426, F_1,27_=20.063, p<0.001). However, there was a much stronger correlation with stride frequency (protraction: y = 0.145 - 0.010x, R^2^=0.691, F_1,27_=60.262, p<0.001; retraction: y = 0.172 - 0.013x, R^2^=0.821, F_1,27_=123.582, p<0.001). The above data shows the duration of protraction is proportional to both speed and stride frequency, but this does little to inform us how the leg is actually moving through space during the swing phase. We now examine variables relating to the position, velocity and acceleration across the duration of the stride.

### Stepping pattern

While stepping pattern diagrams revealed some general trends, there is variation between sequences (Fig. 3). The stepping order for the ipsilateral legs in both Fig. 3B and C is 4 1 3 2. However, at the slowest speed there is no consistent pattern over the sequence (Fig. 3A). The stepping pattern of spiders can be explained by two models of coupling of legs. Considering the coordination of ipsilateral legs, adjacent legs tend to step alternately and adjacent-but-one legs step together. The resulting pattern is a metachronal wave propagating either anteriorly or posteriorly (Fig. 3B). At faster speeds an additional pattern exists, contralateral legs also alternate, resulting in an alternating tetrapod where diagonally adjacent contralateral legs (L1, R2, L3, R4 and R1, L2, R3, L4) step together (Fig. 3C). While movement of the legs is not exactly synchronized, it does appear that appendages can be divided into two functional groups (Fig. 3C(ii)). An alternative representation of leg coordination is a quantitative analysis of relationships between leg pairs, rather than considering several legs at once. This will further illustrate that despite variation between individual steps and trials consistent patterns exist in spider leg coordination.

### Phase relationships between legs

By plotting the time of occurrence of an AEP for a given leg, relative to the stride period of a reference leg, the relationship between timing of leg movements can be scrutinized through a phase histogram (Fig. 4). The mean value of the histogram indicates the relative timing of the movement of the given leg with respect to the reference leg, and the variance gives an indication of the strength of coupling between legs.

**Fig. 4.**
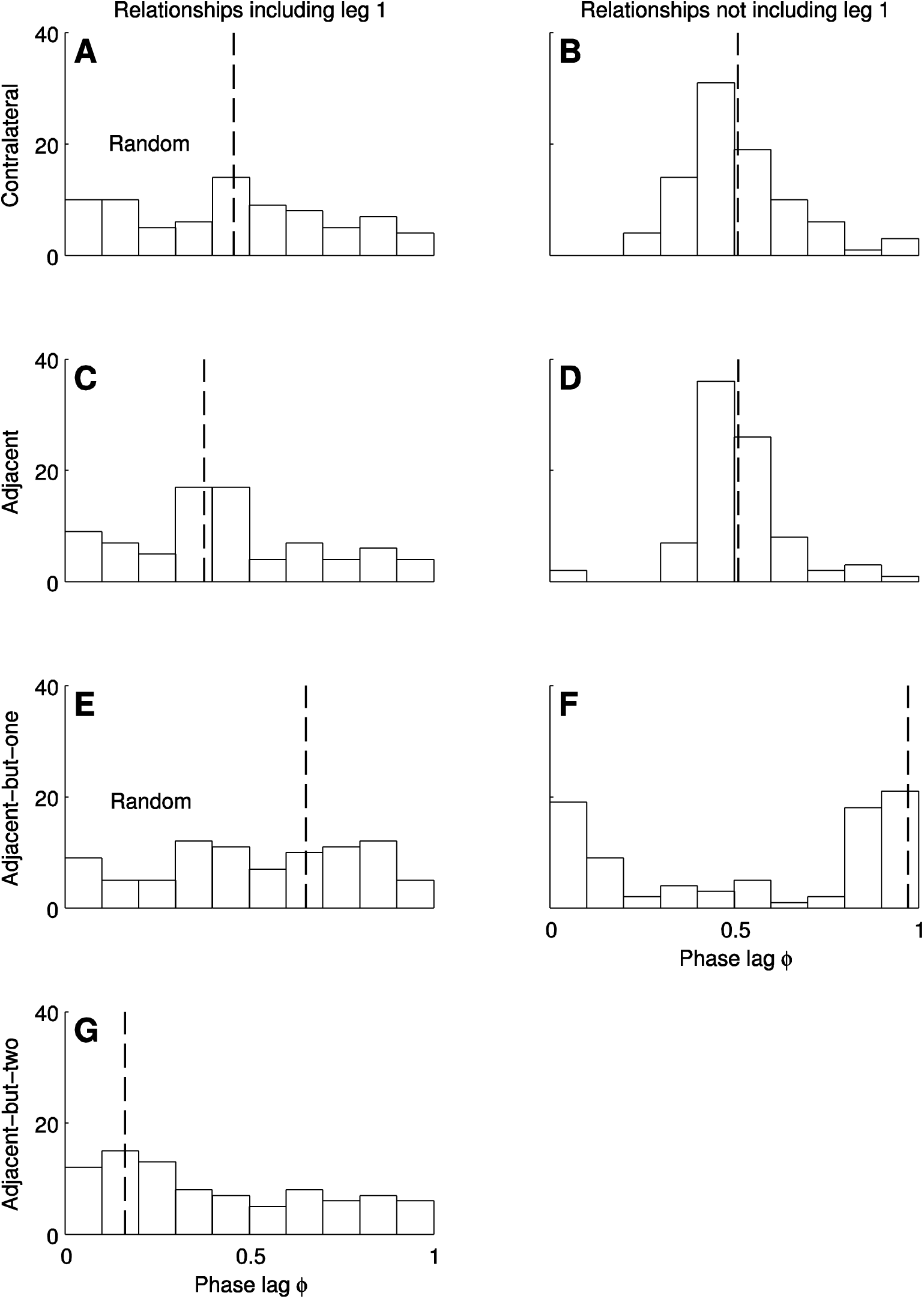
Phase histograms showing the time at which an AEP occurred for a given leg relative within the period of the reference leg. Representative plots were chosen to illustrate different couplings and relationships. (A) Contralateral legs R1:L1. (B) Contralateral legs R4:L4. (C) Adjacent ipsilateral legs R1:R2. (D) Adjacent ipsilateral legs R3:R4. (E) Adjacent-but-one ipsilateral legs R1:R3. (F) Adjacent-but-one ipsilateral legs R2:R4. (G) Adjacent-but-two ipsilateral legs R1:R4. Trends illustrated in C-G were similar for legs on the left-hand side. Dashed lines represent the circular mean. The phase lag distribution for (A) L1:R1 and (E) R1:R3 is random.

The phase lag distributions of contralateral legs all have a mean of approximately 0.5, the value expected from an alternating tetrapod gait (Fig. 4A, B). This result is predicted by Hughes’ (1952) rule for contralateral coupling during normal walking, which states that “each leg alternates with the contralateral limb of the same segment”. In contrast to the idea of alternating sets of legs, the phase relationship in leg pair 1 showed no clear unimodal distribution (p > 0.05, Rayleigh test for uniformity). The phase lags of R1:L1 were rather randomly distributed around the trigonometric circumference (Fig. 4A). All leg pairs have mean phase lags close to 0.5, but appear to have stricter coupling towards the posterior end (mean angle ± circular S.D.: Leg pair 1 = 0.458 ± 0.006, leg pair 2 = 0.498 ± 0.003, leg pair 3 = 0.485 ± 0.002, leg pair 4 = 0.510 ± 0.002).

A similar trend is observed from anterior to posterior in adjacent ipsilateral legs (Fig. 4C, D). Phase lag distributions of pairs containing leg 1 have a greater spread than those which do not contain leg 1 (mean angle ± circular S.D.: L1:L2 = 0.368 ± 0.004, R1:R2 = 0.377 ± 0.005, L2:L3 = 0.437 ± 0.002, R2:R3 = 0.461 ± 0.002, L3:L4 = 0.493 ± 0.002, R3:R4 = 0.512 ± 0.002). It is clear that the mean phase value also increased from the anterior to the posterior pairs of adjacent ipsilateral legs. A possible mechanism for this pattern is provided by Hughes’ (1952) rule for ipsilateral coupling during normal walking, which states that “no fore or middle leg is protracted until the leg behind has taken up its supporting position”.

The remaining couplings for adjacent-but-one and adjacent-but-two ipsilateral legs have probably come about as a result of the above two rules (Fig. 4E-G). Again, results can be split into groupings based on whether the pairing contains leg 1 or not (mean angle ± circular S.D.: L1:L3 = 0.823 ± 0.005, R1:R3 = 0.654 ± 0.005, L2:L4 = 0.936 ± 0.003, R2:R4 = 0.972 ± 0.003, L1:L4 = 0.207 ± 0.005, R1:R4 = 0.163 ± 0.005).

There was no clear unimodal distribution of R1:R3 phase lags (p > 0.05, Rayleigh test for uniformity) and the remaining pairs containing either of the front legs have a more varied distribution than pairs which do not contain either of the front legs. Apart from the pairing of R1:R3 the remaining mean angles are all µ > 0.75 or µ < 0.25 indicating that these legs are moving in-phase.

The possibility that phase lag might change with speed was investigated by angular-linear correlation. The phase lag between left legs 3 and 4 was correlated with both absolute and normalized speed (absolute speed R^2^ = 0.460, p<0.05; normalized speed R^2^ = 0.494, p<0.05). Likewise, the phase lag between left legs 1 and 4 was correlated with both absolute and normalized speed (absolute speed R^2^ = 0.488, p<0.05; normalized speed R^2^ = 0.425, p<0.05). The phase lag of right legs 1 and 2 was correlated with absolute speed but not normalized speed (absolute speed R^2^ = 0.459, p<0.05; normalized speed R^2^ = 0.336, p=0.109).

Mean values from phase lag histograms can be used to reconstruct an average gait (Wilson, 1967). The stepping pattern for ipsilateral legs on the left side is shown in Fig. 5, unsurprisingly the resulting sequence matches the 4 1 3 2 observed in Fig. 3 B and C suggesting that this is the dominant sequence for ipsilateral legs. Likewise, the reconstructed stepping pattern for right ipsilateral legs was also 4 1 3 2, with legs 4 and 2 moving almost simultaneously.

**Fig. 5.**
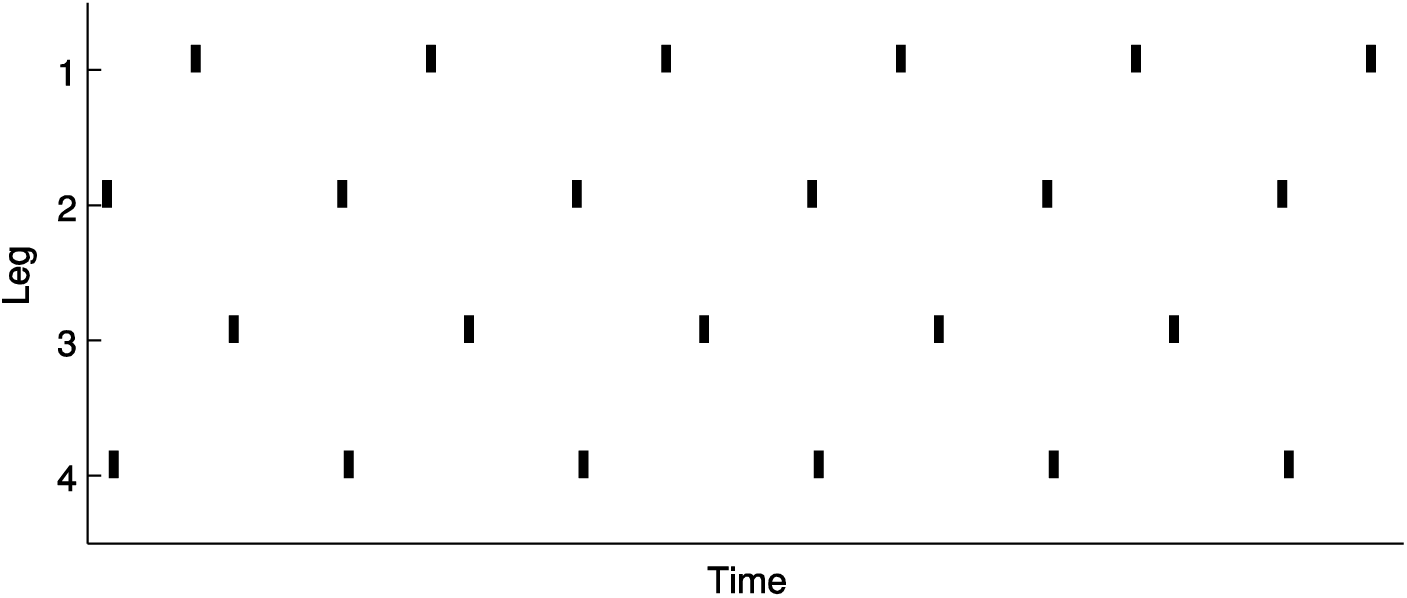
Stepping pattern reconstruction of ipsilateral legs from peak values from the phase histograms. The vertical lines represent the start of each stepping cycle. Reconstructed average gait 4132. Phases used in this case were the left legs; however, using right legs did not alter the reconstructed average gait.

### Position and velocity of legs

Fig. 6 shows the relationship between the angular positions and velocities of legs 1, 2, 3 and 4 for all sequences. This representation of leg movements in their phase plane allows the examination of temporal stability of leg movement. Whilst subjects and sequences both show variation mainly due to differences in speed, these plots prove useful in examining general cyclic patterns of movement which allow us to infer the relative roles of each leg during locomotion.

**Fig. 6.**
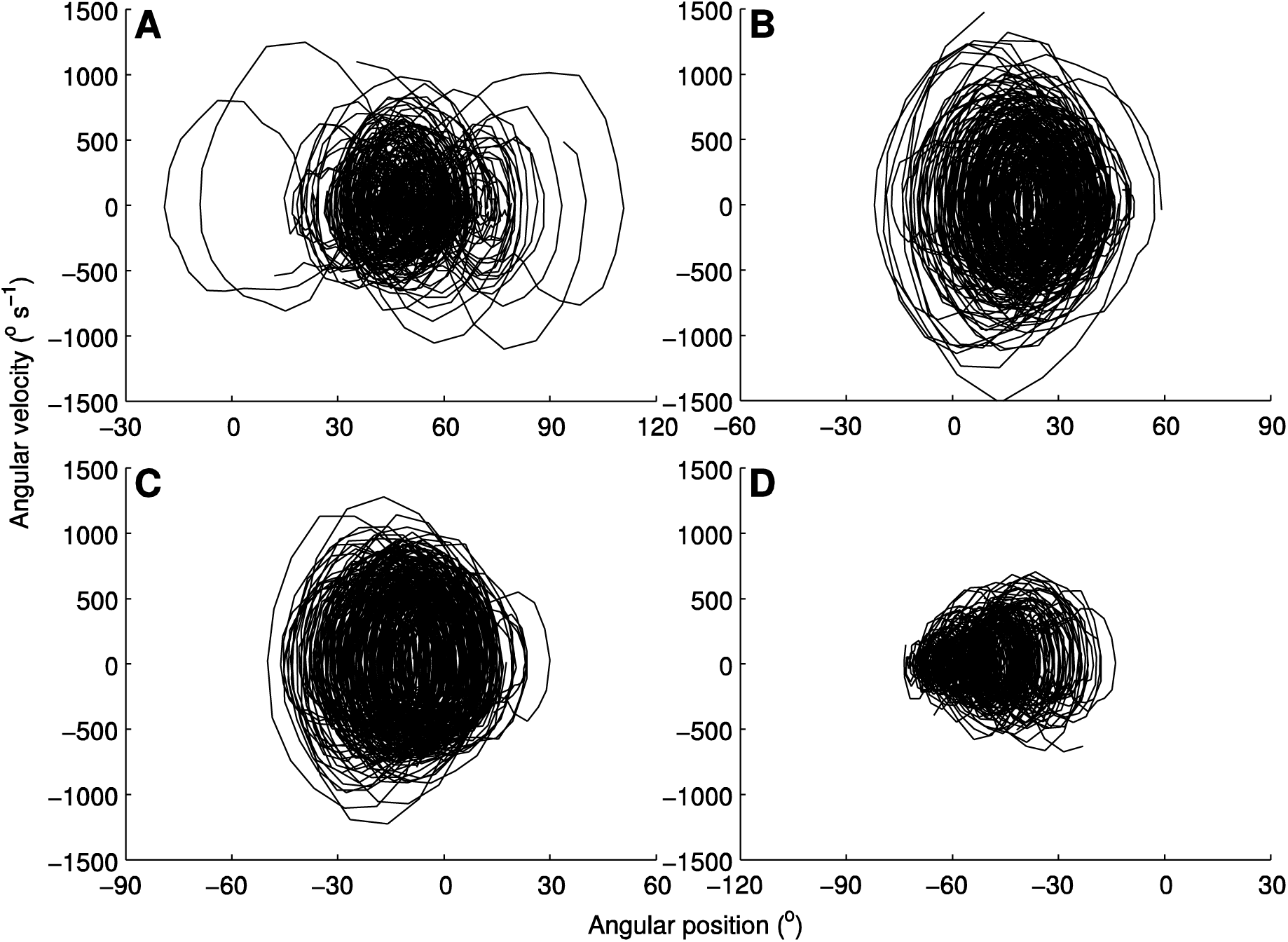
Movements of the legs in their phase plane. Angular speeds plotted as a function of angular positions. Strides occurring during all sequences have been superimposed (N=27). (A) leg 1. (B) leg 2. (C) leg 3. (D) leg 4. Angular position is equal to zero when the leg is perpendicular to the body axis of symmetry, while the x-values differ in magnitude each graph has the same size scale and displays the same range.

The phase portrait of leg 1 (Fig. 6A) again reinforces that the movement of this leg is much more variable than other legs. It should be noted that this variability is not accumulated as a result of combining different sequences but exists in individual sequences. Leg 2 showed reasonably stable cycling which is symmetrically distributed around approximately 20 (Fig. 6B). Likewise, leg 3 showed reasonably stable cycling which is symmetrically distributed around approximately -10 (Fig. 6C). Conversely, leg 4 varied considerably, suggesting that it is possibly acting under the influence of other legs (Fig. 6D).

Average distance covered by the tibia is greater in legs 1 and 4 than legs 2 and 3 (Table 1). Conversely, the angular distance covered in the swing phase by legs 2 and 3 is greater than legs 1 and 4 (Table 1). Likewise, the magnitude of the average angular velocity during the swing phase is larger for legs 2 and 3 than legs 1 and 4 (Table 1). Similarly, during the stance phase legs 2 and 3 experience negative velocities of greater magnitude than legs 1 and 4 (Table 1).

**Table 1.**
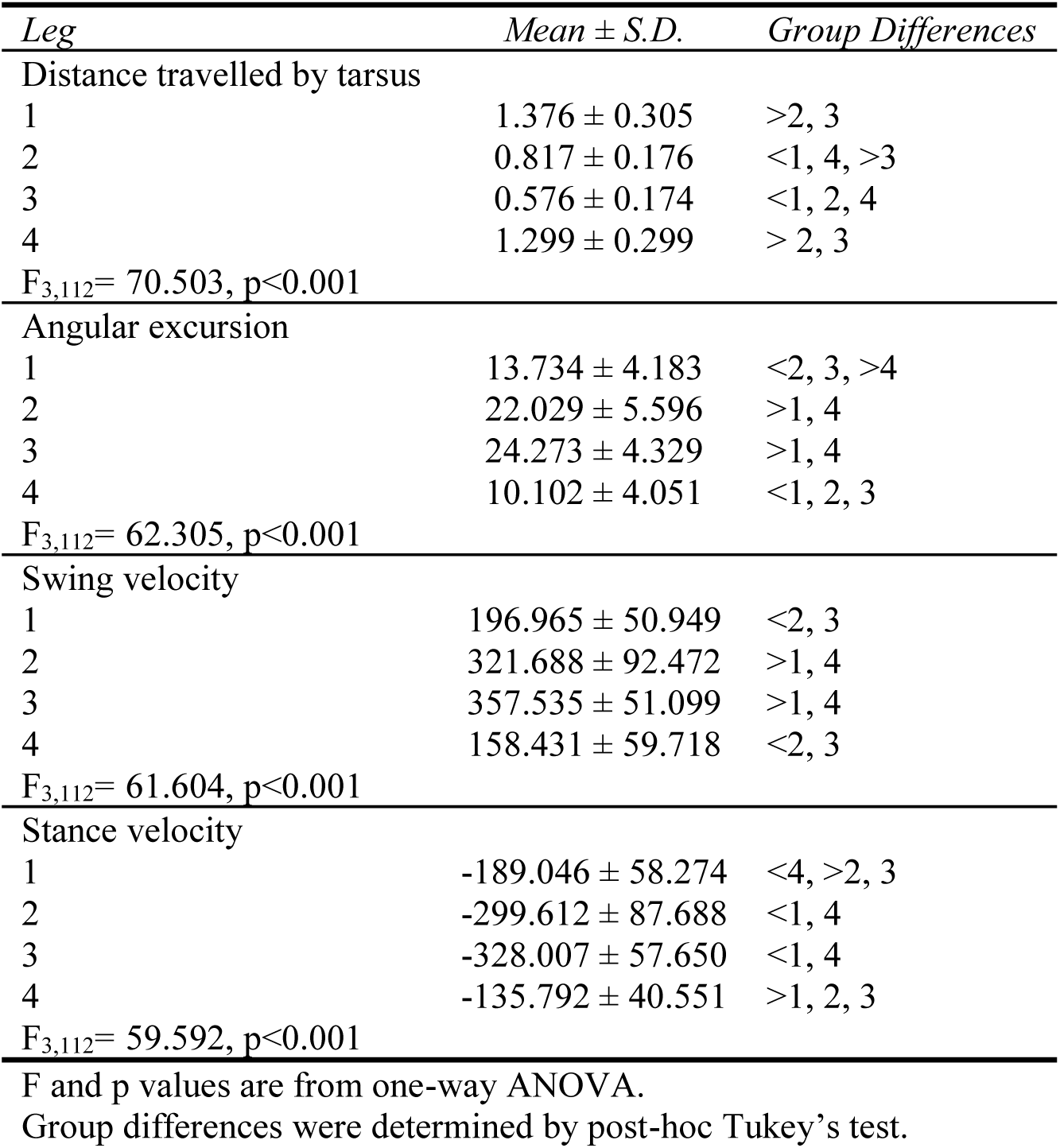
Mean values for distance travelled by the tarsus during swing phase (cm), angular excursion (°), average angular velocity (°s^-1^) in both swing and stance phases for each leg pair.

| *Leg* | *Mean ± S.D.* | *Group Differences* |
| --- | --- | --- |
| Distance travelled by tarsus |  |  |
| 1 | 1.376 ± 0.305 | >2, 3 |
| 2 | 0.817 ± 0.176 | <1, 4, >3 |
| 3 | 0.576 ± 0.174 | <1, 2, 4 |
| 4 | 1.299 ± 0.299 | > 2, 3 |
| $F_{3,112} = 70.503, p < 0.001$ | | |
| Angular excursion |  |  |
| 1 | 13.734 ± 4.183 | <2, 3, >4 |
| 2 | 22.029 ± 5.596 | >1, 4 |
| 3 | 24.273 ± 4.329 | >1, 4 |
| 4 | 10.102 ± 4.051 | <1, 2, 3 |
| $F_{3,112} = 62.305, p < 0.001$ | | |
| Swing velocity |  |  |
| 1 | 196.965 ± 50.949 | <2, 3 |
| 2 | 321.688 ± 92.472 | >1, 4 |
| 3 | 357.535 ± 51.099 | >1, 4 |
| 4 | 158.431 ± 59.718 | <2, 3 |
| $F_{3,112} = 61.604, p < 0.001$ | | |
| Stance velocity |  |  |
| 1 | -189.046 ± 58.274 | <4, >2, 3 |
| 2 | -299.612 ± 87.688 | <1, 4 |
| 3 | -328.007 ± 57.650 | <1, 4 |
| 4 | -135.792 ± 40.551 | >1, 2, 3 |
| $F_{3,112} = 59.592, p < 0.001$ | | |

F and p values are from one-way ANOVA.
Group differences were determined by post-hoc Tukey's test.

### Evaluation of the tracking algorithm

Movies S1-3 included on the in the supplementary material illustrate that the algorithm appears to do a reasonable job of tracking the motion of the spiders’ joints across frames. Generally, motion capture techniques have their accuracy assessed by comparing results to ground truth data. Because we don’t have access to a data set for our study species, we attempt a quantitative evaluation through other techniques below.

Analysing 46 frames where the spider appeared stationary resulted in slightly “jiggly” looking results (see Movie S3). The maximum and minimum position in *x* coordinates differed by 2 pixels and *y* coordinates differed by 1 pixel. Translated in to cm *x* coordinate measurements differed by 0.083 cm and *y* coordinate measurements differed by 0.042 cm. Alternatively, looking at one standard deviation gave 0.557 pixels (0.023 cm) for *x* coordinates and 0.487 pixels (0.020 cm) for *y* coordinates. Individual joint angles were somewhat less accurate than total angle of the leg (Table 2), suggesting that at times the pose estimation fitting results in one angle increasing/decreasing and a joint further down the chain does the opposite.

**Table 2.** Verification of tracking accuracy via two approaches: 1) differences between maximum and minimum values (°) over a sequence of frames where the spider was reasonably stationary; 2) root mean squared differences (°) between forward and reverse tracking for a single sequence.

| *Joint* | Range (°) for stationary frames |  |  |  | Root mean squared differences (°) |  |  |  |
| --- | --- | --- | --- | --- | --- | --- | --- | --- |
|  | Leg |  |  |  | Leg |  |  |  |
|  | 1 | 2 | 3 | 4 | 1 | 2 | 3 | 4 |
| Body-Co | 5<br>(1.502) | 13<br>(3.468) | 6<br>(1.814) | 5<br>(1.268) | 4.739 | 5.395 | 5.185 | 5.827 |
| Co-Tr | 9<br>(2.581) | 14<br>(3.865) | 8<br>(2.465) | 2<br>(0.643) | 6.760 | 5.604 | 7.050 | 5.594 |
| Pa-Ti | 5<br>(1.730) | 14<br>(3.793) | 3<br>(0.643) | 7<br>(2.248) | 6.417 | 3.969 | 4.636 | 3.988 |
| Ti-Me | 9<br>(3.279) | 13<br>(2.899) | 3<br>(0.856) | 3<br>(1.167) | 2.006 | 1.882 | 1.408 | 1.225 |
| Me-Ta | 10<br>(3.528) | 9<br>(2.771) | 13<br>(5.019) | 6<br>(2.464) | 15.851 | 14.035 | 7.689 | 7.521 |
| **Total leg** | 3<br>(0.707) | 3<br>(0.909) | 2<br>(0.662) | 3<br>(0.872) | 6.166 | 4.311 | 3.311 | 3.252 |

Values were averaged over leg pairs.

Comparing the results tracking in one direction and then the reverse direction yielded similar results to above. Position data showed reasonable accuracy with root mean squared difference for position in *x* coordinates of 2.300 pixels and *y* coordinates 0.983 pixels. Translated in to cm *x* coordinate measurements differed by 0.096 cm and *y* coordinate measurements differed by 0.041 cm. Again, the angle for total leg configuration was more accurate than the angles for individual joints (Table 2). Thus, results focused mainly on total leg configuration.

## Discussion

The aim of our study was twofold: to develop a markerless technique for tracking spider locomotion and to use it to characterize the locomotion in *Dolomedes*. While markerless techniques have been an emerging area of human motion capture research for the past couple of decades, they are yet to be widely employed in the area of animal kinematics. We performed detailed analysis of locomotory behaviour in freely moving spiders without attaching physical or painted markers. Our data will now be compared with results obtained previously for spiders and other arthropods; this will be followed by an evaluation of the tracking algorithm.

### Kinematic variables and speed

Both stride length and frequency were directly related to running speed, but stride frequency was more highly correlated with normalized speed and stride length was more highly correlated with absolute speed. This makes sense from a biological perspective because larger animals have longer legs, resulting in longer steps, so if other parameters remain constant this will increase absolute speed. Whereas, when speed is normalized for body length stride length has a reduced role in increasing speed and instead this is achieved by increasing stride frequency. The duration of protraction and retraction were both inversely related to stride frequency and there was no significant difference in regression slopes. This relationship is contrary to the relationship frequently observed in insects, where only retraction decreases with stride frequency and protraction remains constant (reviewed in Wilson, 1966). Results from spider species are conflicting, with some matching our observations (*Trochosa ruricola*, Ward and Humphreys, 1981) and others showing greater similarity to the trends observed in insects (*D. triton*, *Lycosa rabida* - Shultz, 1987; *L. tarantula* - Ward and Humphreys, 1981).

### Inter-leg coordination

The dominant gait exhibited by *D. aquaticus* during locomotion shows similarities to several other spiders that have been studied. Like results from the tarantula (Wilson, 1967) and the jumping spider (Land, 1972), the most common stepping order of ipsilateral legs is 4 1 3 2. The functional grouping of legs is an alternating tetrapod of support, where the odd legs on the left move synchronously with the even legs on the right and vice versa in the second half of the stride. In reality, the functional groups of spiders’ legs do not move in exact synchrony. The reconstructed stepping pattern from phase histograms (Fig. 5) showed legs 2 and 4 starting the stepping cycle almost simultaneously, however leg 3 lagged behind leg 1. Only a small variation in phase can result in similar stepping orders, which also resemble an alternating tetrapod gait (Wilson, 1967). The gaits of spiders *D. triton, L. rabida* (Shultz, 1987), *Hololena adnexa*, *H. curta* (Spagna et al., 2011), *Myrmarachne formicaria* (Shamble et al., 2017) and *T. ruricola* (Ward and Humphreys, 1981) have been interpreted as an alternating. Conversely, some spider species do not approach an alternating gait at even the highest speeds (Moffett and Doell, 1980; Ward and Humphreys, 1981; Weihmann, 2013). It has been argued that rather than a strictly alternating teterapod, spider legs should be treated as two quadrupeds in series, operating slightly out-of-phase with the anterior and posterior quadrupeds less tightly coupled at higher speeds (Biancardi et al., 2011).

Assuming *D. aquaticus* is utilizing an alternating gait, the expected phase lag values for contralateral legs all to be equal to 0.5. Our results found that the coupling of legs 2-4 was approximately equal to 0.5. However, the phase lag distribution for front legs was found to be random. The reader can easily be convinced of this random movement by the examination of the movements of legs in their phase plane (Fig. 6A). This suggests that *D. aquaticus* is actually utilizing its hind three pairs of legs in an alternating tripod gait analogous to insects. The alternating tripod gait predicts that contralateral and adjacent ipsilateral legs are anti-phase (i.e. have phase lags of 0.5) and adjacent-but-one legs are in phase (i.e. have phase lags of 0 or 1). Fig. 4 investigates some of these phase relationships, as hypothesised the phase lags of L2:L3, R2:R3, L3:L4 and R3:R4 have values reasonably close to 0.5; and the phase lags of L2:L4 and R2:R4 have values reasonably close to 1.

Phase lag relationships in the burrow dwelling wolf spider, *L. tarantula*, and the fishing spider, *D. triton*, exhibit similarities to the ones described here. In *L. tarantula* ipsilateral relationships including right leg 1 were random, however anti-phase contralateral coupling between the front legs was maintained (Ward and Humphreys, 1981). Conversely, phase lag values approached 0.5 in adjacent ipsilateral legs in *D. triton*, despite contralateral coupling of the front two pairs of legs being random (Shultz, 1987). *Cupiennius salei* exhibited random contralateral coupling of leg 2, with the remaining contralateral leg pairs phase relations all close to 0.5 (Weihmann, 2013). Ipsilateral relationships further suggested legs may act as two quadrupeds with frontal leg pairs coupled and hind leg pairs coupled and remaining relationships showing or random coupling (Weihmann, 2013). Other species show progressively weaker contralateral coupling in leg pairs from posterior to anterior but distributions were either non-random, or was not verified statistically (*Pardosu tristis* - Moffett and Doell, 1980; *L. rabida* - Shultz, 1987; *Dugesiella hentz* - Wilson, 1967).

The minimum requirement for static stability is three legs in a tripod on the ground at all times, analogous to a stool. Thus, spiders can attain stability with a set of legs to spare, allowing the function of front legs to potentially be modified for functions other than support. The range of results above suggests species are adapted to use front legs for purposes other than support to varying extents. Ward and Humphreys (1981) hypothesised that a possible purpose for these legs is as a sensory apparatus. This was supported by observations by Blickhan and Barth (1985) from tarantulas, which found that the power stroke of the front leg is shorter and the vertical excursion is higher, suggesting that front legs are acting as “feelers” during locomotion. Similarly, the first pair of feet of *Grammostola mollicoma* exhibit an “unusually high trajectory” with longer swing phase, with the authors speculating that they probably play a sensory role (Biancardi et al., 2011). These observations have been suggested as evidence that front legs in spiders have been pre-adapted as an antennae-like structure and this is what allows myrmecomorphic spiders to play such convincing ants, with their front legs raised and moved about like antennae (Reiskind, 1977). However, recent analysis discovered that ant mimics use all eight legs during locomotion which is interrupted with brief pauses where forelegs are raised like ant antennae (Shamble et al., 2017). The sensory role of front legs has been hypothesised in several other arthropods (stick insects - Cruse, 1976; crickets - Harris and Ghiradella, 1980; whip spiders - Santer and Hebets, 2009). While waiting for prey, *D. aquaticus* use the front two pairs of legs to detect vibrations both on the water and land (Forster and Forster, 1999). Many species of spider are nocturnal; in the majority of these species, vision is believed to play a reduced role, or no role at all, in mediating behavioural responses (Foelix, 1982). Thus, it becomes particularly advantageous to use front legs as feelers when hunting at night.

The existence of aerial phases in terrestrial locomotion of two rapidly running spider species, without any obvious gait transitions suggests large, rapidly moving, spiders likely exploit dynamic rather than static stability (Spagna et al., 2011). During passive dynamic walking, there is an impulsive foot-ground collision within each stride, this impact results in loss of energy. The forces involved in the collision are the major determinant of locomotor efficiency (Garcia et al., 2000), thus providing a significant constraint on gait generation. When applied to locomotion in more complex conditions, passive dynamic simulations find it is more efficient to be able to predict ground compliance and pre-emptively modify gait parameters accordingly, rather than recover the trajectory after the collision (Chyou et al., 2011). This implies that if spiders are indeed operating under passive dynamic principles, sensory information would allow disturbances to normal locomotion to be anticipated and compensated for. Therefore, front legs acting as sense organs would play a crucial role in locomotor agility.

Alternatively, the function of these legs has been modified due to habitat usage. Both *D. aquaticus* and *D. triton* are semi-aquatic spiders. During locomotion on the water, a rowing gait involving the legs 2 and 3 is predominately used by *D. triton*, with hind legs sometimes dragged in contact with the water surface (Suter and Wildman, 1999). Lack of use of the front and hind pairs of legs during terrestrial locomotion may be as a result of adaptations relating to the aquatic environment. The fact that *L. tarantula* also exhibits some adaptation in front legs makes this unlikely since this species more or less stays in its burrow waiting for prey to pass (Ward and Humphreys, 1981). Information about terrain height and compliance can be estimated using visual cues. However, this is a non-trivial computational problem requiring eyes looking forward and down, plus neural circuitry capable of performing the analysis. To be effective, especially in dark, highly confined spaces, a spider needs reliable sensing such as that provided by antennae or tactile sensors. Even humans often revert to using poles and probing when on complex unfamiliar terrain. Thus, the most likely explanation is that the front pair of legs is being used as a sensory apparatus.

### Spider leg movements

A previous study, in which walking legs were autotomized, revealed that spiders are capable of adaptive changes and can replace the role of each other during locomotion (Wilson, 1967). Despite this plasticity, the biomechanics of joints are important and our results suggest that each leg plays a distinct role in locomotion. Analysis of the leg trajectories through phase diagrams (Fig. 6) show that each leg exhibits a distinctive pattern. Legs 1 and 2 show mainly positive angular position values, suggesting these legs are used to pull the body forward. Conversely, legs 3 and 4 show mainly negative values so are likely used to push the body. Comparisons between the amplitude of angular velocity of each leg show that legs 2 and 3 undergo much larger positive and negative velocities throughout the cycle. This supports the hypothesis that it is predominantly legs 2 and 3 that are generating propulsive thrust, through pulling and pushing respectively, throughout the locomotory cycle.

The conclusions drawn above are supported by additional experimental results defining leg movement. Analysis of legs during the swing phase (Table 1) found that legs 2 and 3 undergo larger excursions than legs 1 and 4. Additionally, excursion during the swing phase shows a positive relationship with stride duration in legs 3 and 4, suggesting that in order to increase speed these legs decrease the size of excursion and instead increase frequency. Reinforcing the results of the phase diagrams, legs 2 and 3 produced velocities of greater magnitude than legs 1 and 4 in both the swing and stance phases (Table 1). For each leg there was no difference in these magnitudes relative to stride period in the swing phase. However, in the stance phase legs 2 and 3 had negative velocities which were larger in magnitude for shorter stride periods (Table 1).

Specialization in the role of different legs is reasonably common in other arthropods. Jamon and Clarac (1995) found that legs 3 and 4 were the main legs used during crayfish locomotion, although in some animals these patterns were altered so legs 4 and 5 played the dominant role instead. In the rapidly running cockroach, *Periplaneta americana*, different gaits utilize different number of legs generating propulsion. Full and Tu (1991) found that at slow speeds these cockroaches use the alternating tripod gait typical of insect walking, but in order to overcome limitations in stride length, and thus increase speed, animals run primarily on their two hind legs.

### Markerless tracking

The efficiency of motion capture algorithms is generally evaluated on three criteria: the cost and ease of experimental setup, the manual user intervention required and lastly the accuracy of tracking results (Zakotnik and Dürr, 2005). The presented algorithm requires little in terms of equipment, with the key requirements being some sort of behavioural arena which can be filmed from above using a digital video camera. The resulting video sequence is analysed in MATLAB using a constrained kinematic model. The model in this case was rather basic, consisting of a stick spider based on experimentally determined joint lengths and constrained by experimentally determined joint limits (Reußenzehn, 2008). Knowledge of joint limits is not a prerequisite; it does however help constrain the state space. Currently the algorithm requires that the user initializes the size and position of the spider for a given sequence and specify some parameters relating to the CRF. This is done with relative ease through a graphical user interface and is preferable to standard marker-based techniques which may fail in a given frame due to marker occlusion or marker-like reflections etc and require manual correction and re-analysis of subsequent frames. It is somewhat difficult to assess the accuracy of the algorithm without ground truth data; this is attempted through both qualitative and quantitative techniques. Visual inspection video sequences overlaid with the spider model verify that the algorithm performs reasonably well at finding the pose of the subject. For global position, results were highly repeatable with *x-y* coordinates differing by less than 1mm in each dimension when data was compared running the sequence of frames in forward then reverse order. Individual joint angles appeared less accurate than total leg angle (Table 2). Root mean squared differences between forward and reverse tracking for total leg angle average 4.26 across the four legs.

Finally, we openly admit several limitations in the current algorithm and suggest potential improvements. As mentioned previously, individual joints appear somewhat less accurate than the total angle of the leg, suggesting that the optimization framework is causing one angle to increase/decrease and a joint further down the limb to do the opposite. An attempt was made to avoid this by having angular rotations occur from segments closest to the body outwards, but because the angles and corresponding energy term are recalculated several times per frame this was not as successful as anticipated. If improvements are made to the speed of the algorithm, it becomes feasible to run each sequence a number of times and then calculate pose based on the distribution of results rather than a single run. Similarly, repeated analysis could be done on each frame and a disambiguation method could be used to select one of the poses as the correct solution. Such a technique has been implemented for marker-based motion capture which uses stochastic optimization, rather than relying solely on segmentation, to determine the position of markers (Zakotnik and Dürr, 2005). The disambiguation method described in this research is based on the Viterbi method for Markov chains and works by processing the entire sequence, then selecting the trajectory which is globally smoothest for all joint angles and frames (Zakotnik and Dürr, 2005).

Knowledge about the mechanical properties of spider limbs and the physics involved in locomotion helps constrain the state space required for movement. The more we know about the spiders’ mechanical design, the easier the task of tracking becomes. The current tracking algorithm only makes use of the very basic assumption of constant velocity to predict the pose of the spider in the next frame. An accurate physical model of the spider would improve these predictions and make the statistical modelling easier.

With monocular techniques, many possible poses can explain the same spider shape observed in the image. Multiple views can be used to reduce the ambiguity in pose. To maintain the simplicity of a single camera setup, the view from the top and side can be simultaneously recorded by arranging a mirror set at 45 from the animal plane. Using the PoseCut approach multiple views can be easily incorporated into a single optimization framework (Bray et al., 2006).

### Conclusions

The markerless tracking algorithm proved successful in tracking spiders during locomotion. This represents a step away from the traditional marker-based techniques towards computer vision-based approaches, which helps avoid some of the difficulties of marking small animals.

Stride frequency showed a greater correlation with normalized speed than absolute speed and stride length showed a greater correlation with absolute speed than normalized speed. This suggests that larger animals increase their speed by increasing stride length, whereas it is possible for smaller animals to increase their speed by increasing stride frequency. Contrary to the relationship frequently observed in insects and some spiders, both protraction and retraction periods decreased with speed.

Each leg was found to contribute in a specific manner to locomotion. Movements of front legs were random suggesting they play some other role, possibly sensory, rather than contributing to stability. The 2nd and 3rd pairs appeared to play a more dominant role in generating propulsive force, with hind legs probably contributing more to stability than propulsion.

Now that there is a basic understanding of the biomechanics behind terrestrial locomotion, it is possible to begin exploring how locomotion varies under different conditions and how the nervous system produces these changes.

**List of abbreviations**
A: Anterior
AEP: Anterior extreme position
CRF: Conditional random field
GUI: Graphical user interface L Left
P: Posterior
PEP: Posterior extreme position
R: Right

## Acknowledgements

The authors would like to thank to Rasmas Jensen, who made his human tracking code readily available. Ken Miller and Murray McKenzie who provided technical assistance and Dr Phil Bishop who offered spider advice. Stefan Reußenzehn provided the digitized surface points of *Dolomedes* body and legs and joint angle limits that were used to construct the spider model.

## Author contributions

Conceptualization: K.F.P., M.G.P.; Methodology: K.F.P., M.G.P.; Software: K.F.P.; Validation: K.F.P.; Formal analysis: K.F.P.; Investigation: K.F.P.; Resources: K.F.P., M.G.P.; Data curation: K.F.P.; Writing - original draft: K.F.P..; Writing - review & editing K.F.P., M.G.P.; Visualization: K.F.P.; Supervision: M.G.P.; Project administration: K.F.P., M.G.P., Funding acquisition: M.G.P.

